# *In vivo* characterization of chick embryo mesoderm by optical coherence tomography assisted microindentation

**DOI:** 10.1101/2020.03.10.985028

**Authors:** Marica Marrese, Nelda Antonovaité, Ben K.A. Nelemans, Ariana Ahmadzada, Davide Iannuzzi, Theodoor H. Smit

## Abstract

Embryos are growing organisms with highly heterogeneous properties in space and time. Understanding the mechanical properties is a crucial prerequisite for the investigation of morphogenesis. During the last ten years, new techniques have been developed to evaluate the mechanical properties of biological tissues *in vivo*. To address this need, we employed a new instrument that, via the combination of micro-indentation with Optical Coherence Tomography (OCT), allows us to determine both, the spatial distribution of mechanical properties of chick embryos and the structural changes in real-time provided by OCT. We report here the stiffness measurements on live chicken mesoderm during somite formation, from the mesenchymal tailbud to the epithelialized somites. The storage modulus of the mesoderm increases from (176±18) Pa in the tail up to (716±117) Pa in the somitic region. The midline has a storage modulus of (947±111) Pa in the caudal presomitic mesoderm, indicating a stiff rod along the body axis, which thereby mechanically supports the surrounding tissue. The difference in stiffness between midline and presomitic mesoderm decreases as the mesoderm forms somites. The viscoelastic response of the somites develops further until somite IV, which is commensurate with the slow process of epithelization of somites between S0 and SIV.

Overall, this study provides an efficient method for the biomechanical characterization of soft biological tissues *in vivo* and shows that the mechanical properties strongly relate to different morphological features of the investigated regions.

## Introduction

Morphogenesis is a continuous process of cell migration, tissue deformation, and growth. It is a self-organized patterning process orchestrated by the properties of the cells, which are controlled by gene expressions and chemical and physical signaling. While biochemical signals are known to play a fundamental role in the control of tissue morphogenesis (Alan Mathison Turing, 1952; Gjorevski and Nelson, 2010; Miller and Davidson, 2013; Pourquié, 2003), several *in vitro* and *in vivo* studies (Chevalier et al., 2016; Discher et al., 2005; Engler et al., 2006; Filas et al., 2015) have shown the relevance of mechanical cues in the control of cell behavior that are central for the developmental processes. Unraveling the functional role of mechanical forces in morphogenesis is, therefore, a crucial research topic for the developmental biologists. Specifically, the processes during somite formation along the rostrocaudal axis of the embryo, such as the changing of extracellular matrix (ECM) composition, the differential migration and the active cell contraction of epithelial cells of the mesoderm, suggest that there should be differences in mechanical properties along the rostrocaudal axis of the embryo. However, the lack of methodologies enabling precise and quantitative measurements of mechanical properties of live tissues has hindered an exhaustive understanding of the role of mechanics in embryonic development. In our earlier work (Marrese et al., 2019), we proposed an experimental platform that combines micro-indentation and optical coherence tomography to assess mechanical properties in paraformaldehyde-fixed embryos. There, we have demonstrated a relation between local mechanical properties and tissue morphology for three main embryonic regions of interest: the tail, the presomitic mesoderm, and the somitic mesoderm. Although in our previous study, we reported a stiffness map on the somitic region for live embryos, we investigate here the viscoelastic properties of the entire live chicken embryo mesoderm during somite formation. To that end, HH9-HH11 chicken embryos were cultured in filter paper sandwiches, immobilized in agarose, and indented along the embryo with the ferrule-top indenter, while the structure was locally imaged via optical coherence tomography (OCT). The simultaneous use of these two technologies allows one to perform systematic studies on two interconnected topics: on the one hand, the mechanical properties of the embryos that can be characterized through tissue microindentation; on the other hand, and the change in shape that occurs during morphogenesis. Therefore, we present a local mechanical characterization of live embryos that extends our previous work (Marrese et al., 2019) by highlighting the mechanical heterogeneity and the strong viscoelastic nature of the embryonic tissue *in vivo*. We further demonstrate that, while there are substantial differences in absolute viscoelastic responses between individual embryos, the relative trends among anatomical regions are similar and reasonably related to the maturation of the presomitic mesoderm (PSM) and midline in the trunk and tail. This study opens new avenues to explore how mechanics can contribute to shaping embryonic tissues and how it affects cell behavior within developing embryos.

## Results and Discussion

The indenter is based on ferrule-top technology (Chavan et al., 2010; Gruca et al., 2010), where a micro-machined cantilever, operating as a force transducer and equipped with a spherical tip, is used to determine the viscoelastic properties of the embryo via depth-controlled oscillatory ramp indentation profile (Antonovaite et al., 2018; Marrese et al., 2019; Van Hoorn et al., 2016). The OCT system images the embryonic structures during the indentation measurements and allows localization of the indentation points and evaluation of the quality and the immobilization of the sample. The details of the experimental setup (Fig. 1) and sample preparation are briefly reviewed in the Method section and fully reported elsewhere (Marrese et al., 2019).

**Figure 1.**
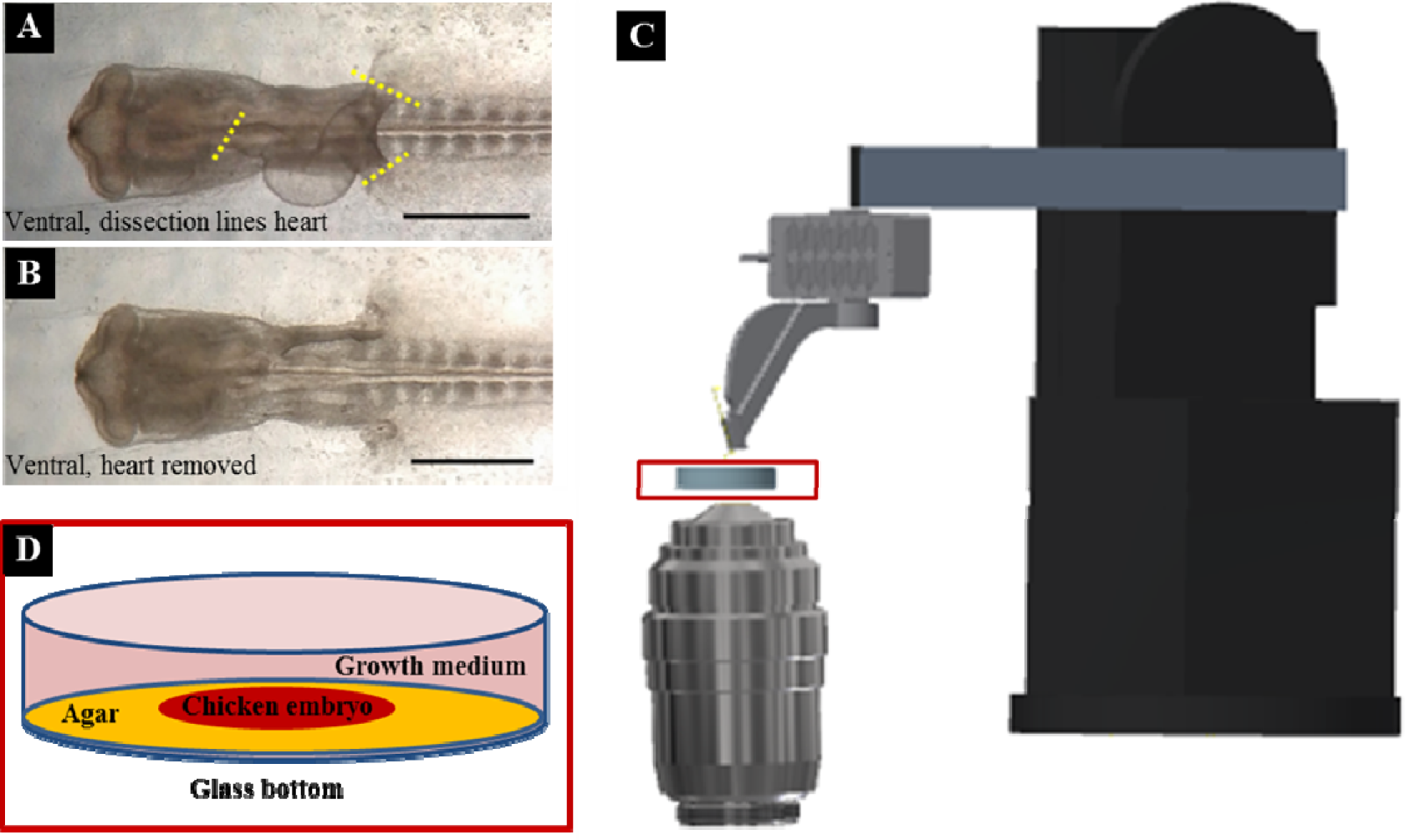
Schematic view of the setup and the sample preparation. (A) Ventral view of an HH11 chicken embryo (40hpf). Yellow lines show the dissection sites to remove the embryonic heart tube, to prevent the beating heart from disturbing the measurements. (B) The same embryo as in (A), after dissection of its heart tube. (C) A ferrule-top probe is equipped with an optical fiber for interferometric readout of the cantilever and with a spherical tip to indent the sample. The probe is mounted on the Z-piezoelectric actuator, which is solidly attached to an XYZ manipulator. The OCT is employed in inverted mode. (D) The embryo is embedded in agarose on its dorsal side, while the ventral side is approachable for measurements and immersed in the growth medium.

Performing a full indentation map at 50 µm resolution along the embryo with the proposed depth-controlled oscillatory profile is time-consuming. A single indentation takes ∼60 s while a new somite is formed every 80 minutes; thus, it is not feasible to map the entire embryo at the same developmental stage *in vivo.* Therefore, to preserve the spatial accuracy along the embryo, we limited indentations to eight lines along the rostrocaudal axis of the mesoderm that are anatomically different. For further details regarding the investigated areas, we refer the reader to Fig. 2 and 3.

**Figure 2.**
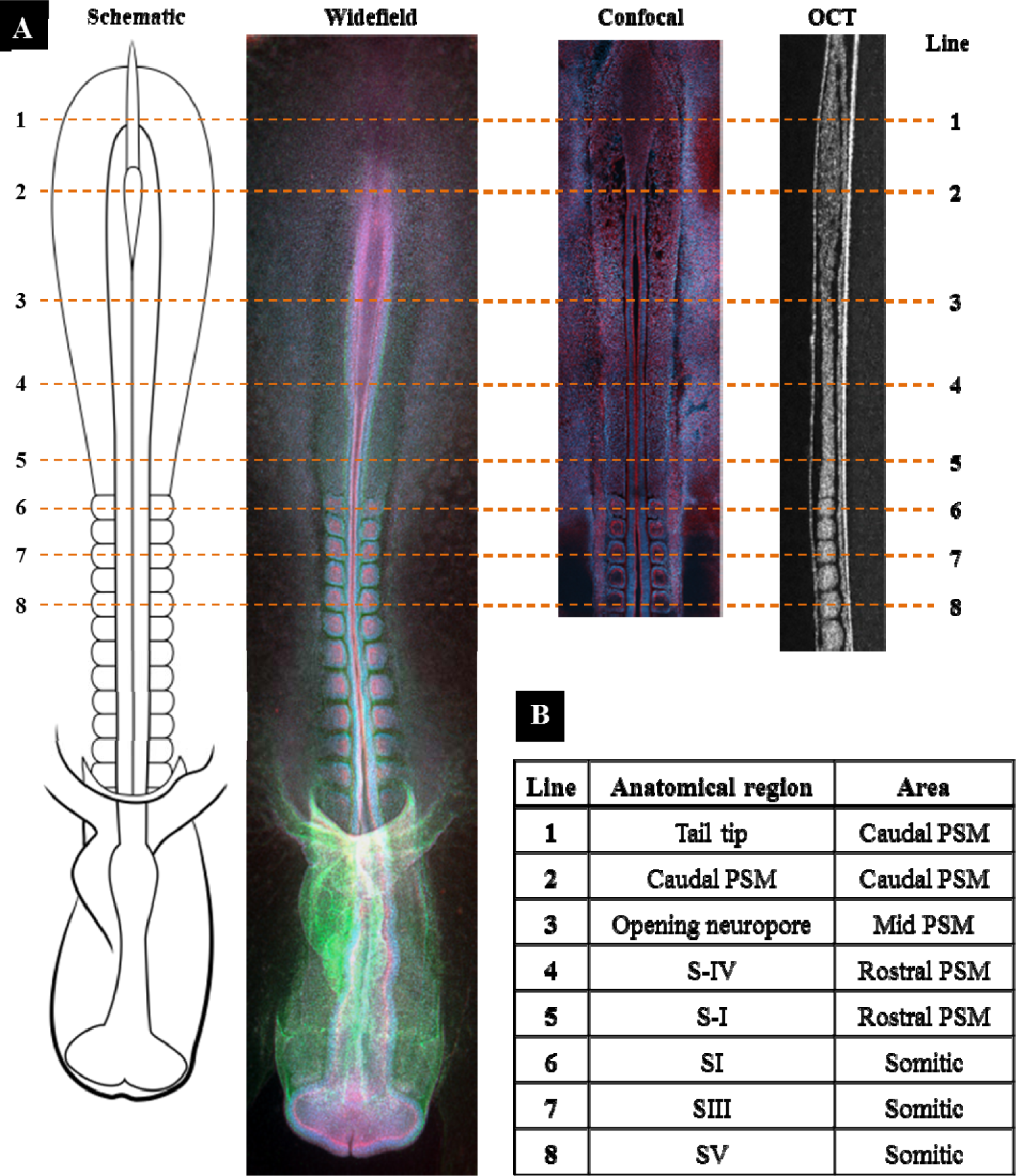
Sagittal embryo indentation points. (A) Embryos were indented with eight transverse lines, at 10 positions with 50 µm steps, across the rostrocaudal axis while visualized by OCT. The lines are shown imposed on a schematic embryo and a widefield immunograph. Next, are a parasagittal confocal section and a midsagittal OCT section, through the mesoderm. Rostral is up, and caudal is down. Immunostainings are red (actin), green (fibronectin), blue (nuclei). The scale bar is 500 µm for all images. (B) Anatomical regions that were indented, from rostral to caudal. The scale bar is 100 µm.

**Figure 3.**
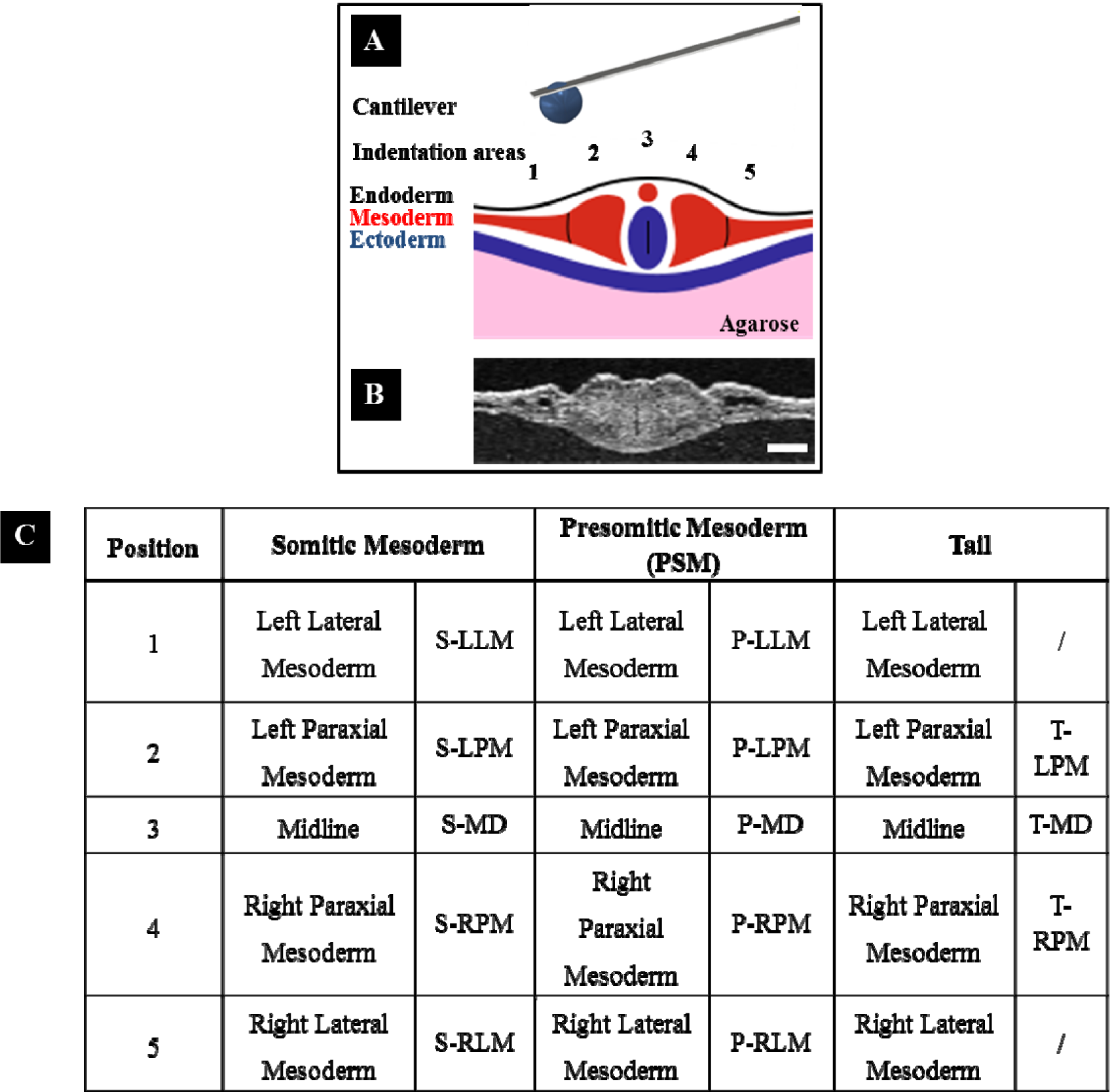
Transversal embryo indentation points. (A) Schematic view and (B) OCT cross-section of the indentation positions of an HH11 embryo embedded in agarose. The scale bar is 100 µm. (C) Anatomical regions that were indented across the embryo. Abbreviations of the region of interest from left to right: (LLM) left lateral mesoderm, (LPM) left paraxial mesoderm, (MD) midline, (RPM) right paraxial mesoderm, (RLM) right lateral mesoderm.

Fig. 4 shows the averaged storage modulus (***E***′) and loss modulus (***E***″) of 12 *in vivo* HH9-HH11 chicken embryos obtained for eight positions along the rostrocaudal axis of the mesoderm. The measurements were done at an averaged strain of ∼8% and 2.5 Hz oscillation frequency. For each of the eight positions along the embryo, the plot shows the distribution of ***E***′ and ***E***″ for five regions of interest: left and right lateral mesoderm, left and right paraxial mesoderm, and midline. From the data in Fig. 4 along with OCT images, one can observe a consistent correspondence between the morphology of the indented regions and their mechanical properties. The stiffness difference between the paraxial mesoderm and the midline is more significant in the PSM and the tail than in the somitic area. In the caudal PSM, the paraxial mesoderm is very soft, while the midline stiffness ***E***′ significantly increases from (270±36) Pa caudally to (947±111) Pa more cranially (Fig. 4, lines 8 to 6, mean ± SEM, p=0.0009; 0.005, Wilcoxon rank-sum test; Fig. S1). At the somitic levels, the midline is still the stiffest structure, but the difference with the stiffness of the somites is negligible (Fig. 4, lines 1, 2, 3, p=1, 0.89, 0.65 left side and p=0.25, 0.27, 0.39 right side, Wilcoxon rank-sum test).

**Figure 4.**
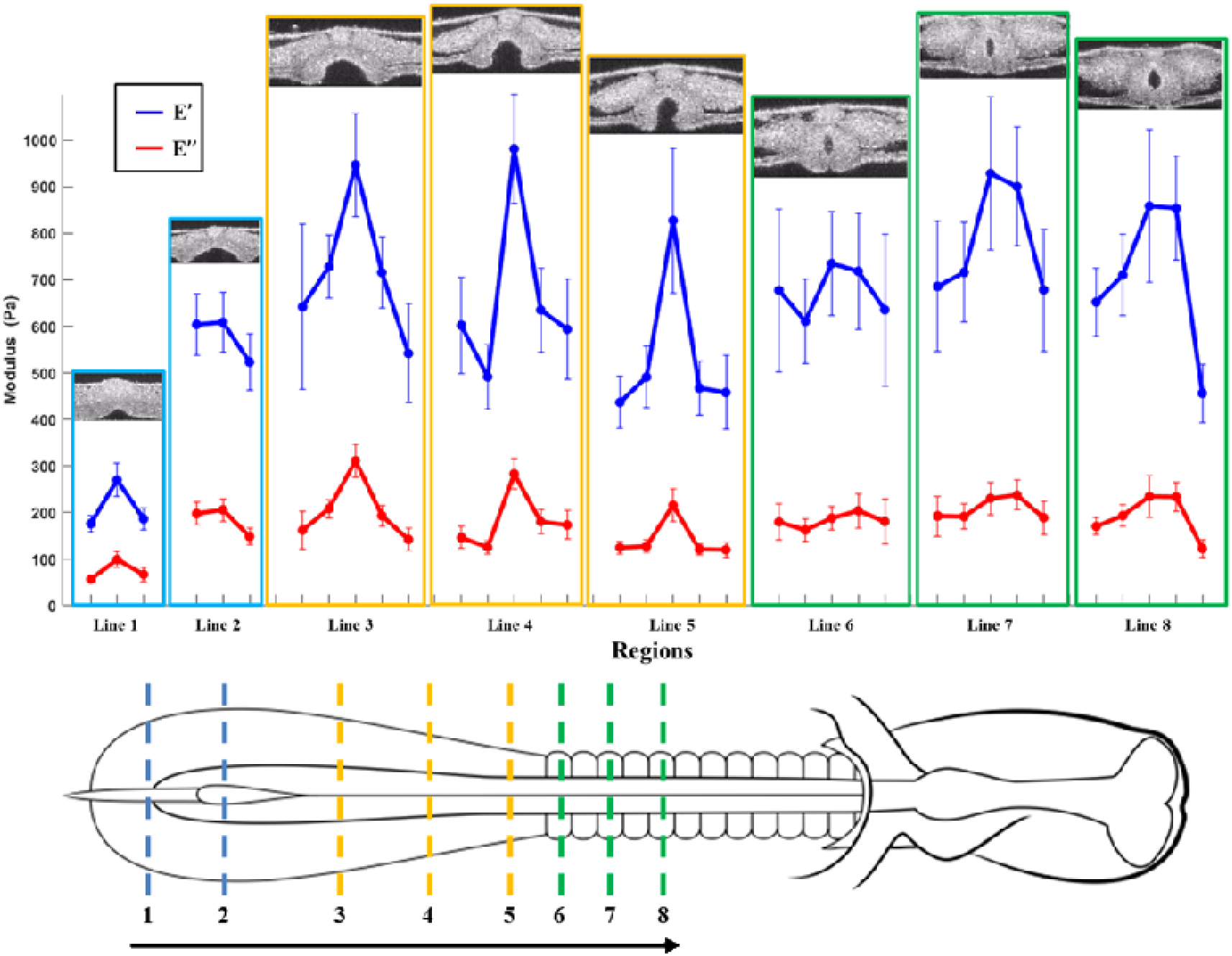
Averaged storage (, blue) and loss (, red) modulus along the embryo. Transverse OCT sections show the positions of line 1 (most rostral) to line 8 (most caudal). Data points are averages of 12 embryos, with SEM error bars. Every line shows five regions of interest, from left to right, these are: (1) right lateral mesoderm, (2) right paraxial mesoderm, (3) midline, (4) left paraxial mesoderm, (5) left lateral mesoderm. The black arrow indicates the locations of the eight lines from tail to somites.

Somites III to V are slightly stiffer than somite I, but not significantly (Fig. 4 lines 1 and 2 vs. 3; p=0.18, 0.17 left side, and p=0.66, 0.07 right side, Wilcoxon rank-sum test). Similarly, the paraxial mesoderm increases its storage modulus on average from (527±38) Pa in the rostral PSM up to (746±44) Pa in the somitic region (Fig. 4 lines 1, 2, 3 vs. 4 and 5: p=0.006, Wilcoxon rank-sum test).

Furthermore, Fig. 4 shows a significant variation in stiffness in the caudal PSM when compared with the tail for both the midline (609±63 and 270±35 Pa, respectively, p=0.0002) and the paraxial mesoderm (558±44 and 181±14 Pa, p=0.0001, respectively, Wilcoxon rank-sum test).

The observed trends can be logically related to the maturation of the chicken embryo (see micrographs in Fig. 2). The caudal PSM is characterized, in fact, by stem cell-like mesenchymal cells that migrate actively with large intercellular space and lack a mature extracellular matrix (ECM) (Fig. 2A, confocal, and OCT section) (Bénazéraf et al., 2010). Gradually, fibronectin and laminin become more abundant and interconnect rostrally (Fig. 2A widefield). This aids in anchoring the PSM cells by providing a substrate on which they can undergo collective migration and mesenchymal-epithelial transition (MET) to form epithelial spheres(Cheney and Lash, 1984; Oster et al., 1983; Sato et al., 2017). The PSM cells compact together, adhere to the ECM and each other, and become more contractile (Fig. 2A confocal section, compare caudal PSM with rostral PSM) (Bard, 1988; Goto, 2012), thereby promoting fibronectin assembly. Concurrently, the notochord and neural plate quickly develop a high stiffness (Fig. 2, lines 8 to 6). This behavior seems to support the idea that the notochord is not only an organizer center for chemical signaling but also acts as an ‘embryonic spine’ that plays a significant role in the mechanical integrity of the early embryo (Corallo et al., 2015). Next, the neural plate rostrally folds into the neural tube, and this morphogenetic movement could be due to a stiffer tissue that undergoes neurulation (Fig. 3, lines 5 and 6). This finding agrees with previous studies on the Xenopus, where morphogenetic transformations are preceded by stiffening of the structures (Zhou et al., 2009). After neurulation, the neural tube keeps developing, but the presence of the lumen in the tube could lead to a softer tissue able to deform more when indented if compared to the compact neural groove (Fig. 4, lines 4, 5 vs. 1, 2, 3; Fig. S2).

Dynamic indentation reveals that a viscous component is present in embryonic tissues as well (with *E*′ ∼3*E*″); this is illustrated by the values of loss modulus *E*’’ in Fig. 4. To describe the energy damping potential of the embryo under loading, the averaged damping factor, tan(*φ*), is shown in Fig. 5 as the ratio between loss and storage modulus (*E*’’/*E*’). The values of tan(*φ*) are comparable for the paraxial mesoderm, the midline, and the lateral mesoderm for the somitic area (p=0.27-0.97). However, in the tail and PSM, damping capability is higher for midline and lower for paraxial mesoderm (p=0.0001, 0.02; p=0.02, 0.09, left and right, respectively, Wilcoxon rank-sum test) with the tail having overall highest damping factor (tan(*φ*)=0.30-0.33 vs. tan(*φ*)=0.26-0.29). Specifically, this finding could be related to the status of development with the epithelial tissues (more mature and with more extracellular matrix) being more elastic and less viscous due to their nature.

**Figure 5.**
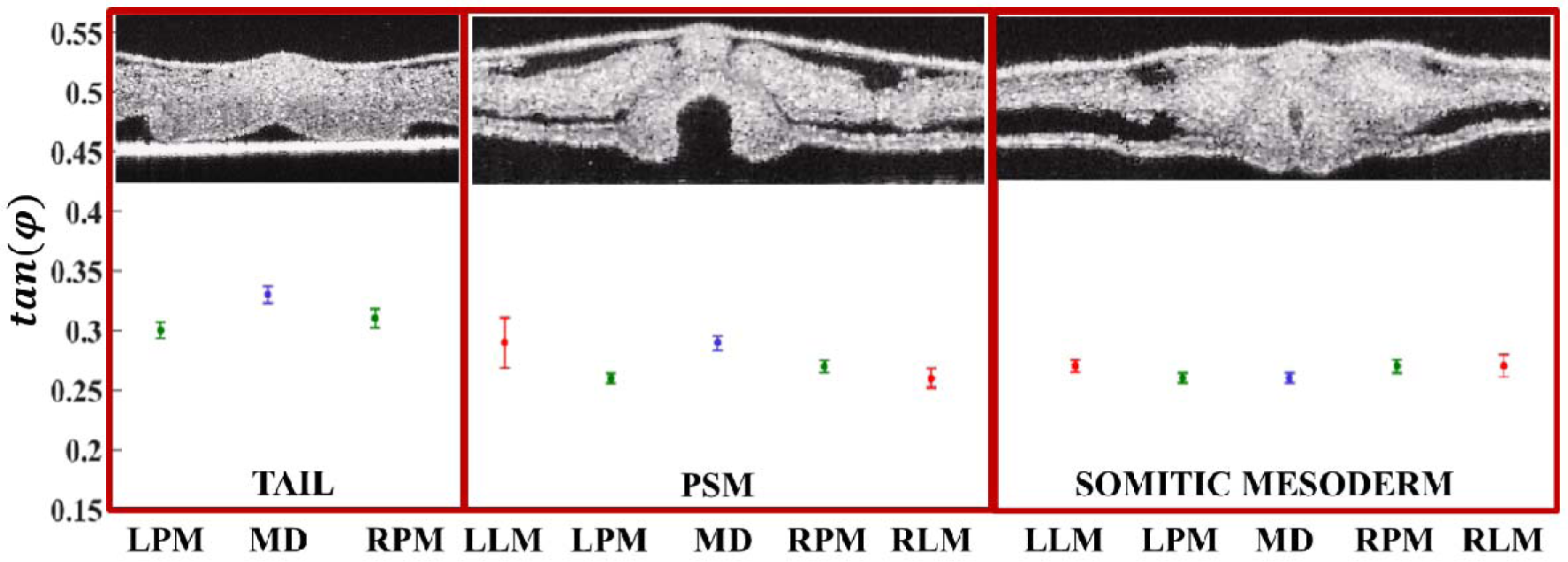
Averaged damping factor of 12 embryos over lines from the same areas (somatic and presomitic mesoderm and tail) with SEM bars. Abbreviations of the region of interest from left to right: (LLM) left lateral mesoderm, (LPM) left paraxial mesoderm, (MD) midline, (RPM) right paraxial mesoderm, (RLM) right lateral mesoderm.

It is worth to mention that averaging the viscoelasticity values from the same rostrocaudal regions over embryos resulted in logical trends of stiffness, that are commensurate with epithelization and matrix formation. Nevertheless, we observed substantial variation in viscoelasticity between single embryos. It appears that the biomechanical properties of the embryonic tissues may vary with age and quality of the embryo. As a point in case, differences in the handling of the embryos could have influenced their viability and, thus, their mechanical properties. Furthermore, the indentations are influenced by the accurateness of positioning the probe tip: small variations in positioning the probe along the embryonic structures could have led to local shearing or slipping of the probe along the tissue.

It is interesting to note that some of the mechanical features observed *in vivo* are different from the ones described for the fixed embryo (Marrese et al. 2019). The mechanical maps reported previously for the formaldehyde-fixed embryo (Marrese et al. 2019) showed an increase of stiffness along the mesoderm from the caudal tip to the rostral somites, possibly related to the effect of the formaldehyde to fix tissue by cross-linking of the biopolymers. This result is not confirmed for the live embryos, where the midline stiffness is already high in the tailbud. This finding shows the effect of formaldehyde fixation on two complex embryonic structures: the notochord and the neural tube. By measuring *in vivo*, our instrument seems to be able to sense how the PSM (Fig. 4, lines 5 and 6) is characterized by the opening of the neuropore, which has a large cell contraction as it closes to form a tube. Moreover, for the *in vivo* embryo, the low stiffness in the tail region is more evident if compared to the formaldehyde-fixed embryos, possibly due to a lack of structural components such as the neural tube and the notochord. One can argue/state that *in vivo*, we are able to sense mechanical properties caused by active biomechanical processes, such as stiffening by cellular contraction, while the measurements on fixed embryo are strictly linked to tissue morphology. This behavior seems to indicate that chemical fixation has two effects on the live soft tissue: it increases tissue stiffness and reduces the damping properties of the embryonic tissues.

Comparing elasticity of the midline, the paraxial and lateral mesoderm before and after fixation, the average storage and loss moduli were found to be a factor ∼2 times higher after fixation. In addition, the trends of tan(*φ*) differs from the results obtained for the paraformaldehyde-fixed embryos. Furthermore, tan(*φ*) is overall lower for the live embryo (∼1.4 times). Moreover, it is interesting to mention that while observing the morphological features of the embryonic structure *in vivo* and after 2 hours fixation via OCT, some regions of the embryo appeared to be structurally different: the morphology of the embryo seems, in fact, more compact and dense (Fig. S.3). By taking a closer look at the OCT images in fig.S.3 for each of the analyzed location, one can speculate that the tissue after fixation becomes denser and contains less fluid and, thus, the loss modulus increases more than storage modulus resulting in a higher tan(*φ*). These findings provide key insights into differences between *in vivo* and chemically treated tissue and underline the importance of using *in vivo* tissue to study the biomechanics of embryos.

Specifically, our measurements show that the midline already stiffens near the tail and essentially acts as an embryonic spine. The damping factor is reduced when moving from tail to head, indicating a more elastic behavior for the more mature embryonic structures. Lastly, the method allows for sensitive detection of structurally distinct embryonic areas, both visually and mechanically. We demonstrate that our platform can reliably measure the viscoelastic properties of the tissue with more precision than previous studies (Agero et al., 2010), and allows one to discriminate between the small embryonic structures like somites and neural tube. Finally, since mechanical stress can modulate physiological processes at the cellular and tissue level, we expect that this study will support a significant step forward in gaining new insights on the relationship between altered morphogenesis, stiffness, and pathologies during the embryonic morphogenesis.

## Materials and Methods

### Chicken embryo cultures

The embryo cultures were prepared as described somewhere else (Marrese et al., 2019). Briefly, fertilized chicken eggs, white leghorns, *Gallus gallus domesticus* (Linnaeus, 1758), were obtained from Drost B.V. (Loosdrecht, The Netherlands), incubated at 37.5 °C in a moist atmosphere, and automatically turned every hour. After incubation for approximately 41h, HH9-HH11 chicken embryos (Hamburger and Hamilton, 1992) were explanted using filter paper carriers (Chapman et al., 2001) cultured *ex ovo* as modified submerged filter paper sandwiches (Schmitz et al., 2016), immobilized in agarose and immersed in growth medium (Palmeirim et al., 1997).

To avoid disturbance of the measurements by the beating of the heart that might develop, the heart tube of the ventral side of sandwiched embryos was removed (Fig. 1A and B). This does not appear to inhibit further development of the spinal structures in the chick embryo. To prevent dehydration of the live embryo during the LGT agarose curing, a droplet of the medium was carefully brought on top of the embryo, without touching the curing agarose. After approximately 3 minutes, the culture was placed in the indentation box, submerged in 25 ml of the growth medium and anatomically aligned under the OCT to precisely discriminate each indentation location.

The growth medium consisted of medium 199 GlutaMax (Invitrogen, ref. 41150-020; 4ºC), 10% chicken serum (GIBCO, ref. 16110-082; −20ºC), 5% dialyzed fetal bovine serum (FBS) (GIBCO ref. 26400-036; −20ºC) and 1% of a 10000 U/ml stock solution of Penicillin/Streptomycin (GIBCO ref. 15140-122; −20ºC).

### Experimental setup

The setup consists of a cantilever-based indentation arm, an OCT imaging system, and a sample holder (Fig. 1). The indenter is based on a micro-machined cantilever, operating as a force transducer. An extensive description and validation of the experimental setup have been reported in our previous publication (Marrese et al., 2019). Briefly, for indentation measurements on live embryos, cantilevers with spring constant in the range of 0.34-1.2 N/m, and spheres radius between 54 and 69 µm were used. Indentations were performed in a depth-controlled mode by using an oscillatory ramp profile at indentation speed of ∼0.5 µm/s, maximum indentation depth of 30 µm, and amplitude and frequency of oscillations were 0.25 µm and 2.5 Hz, respectively. Load-indentation data were used to extract storage and loss moduli. Storage modulus values at an averaged strain of 8±1% (corresponding to a depth of ∼10-12 µm) were selected for regional comparisons, accomplishing the requirements of *h* < 10% of the sample thickness and small strain approximation (Lin et al., 2009).

To find anatomical locations and follow each indentation experiment, the embryos were scanned with a spectral-domain SD-OCT (Telesto II series, Thorlabs GmbH, Germany) in inverted mode, as reported elsewhere (Marrese et al., 2019).

To perform a full mechanical characterization of the embryonic tissues, we focus on eight positions along the rostrocaudal axis of the mesoderm that show anatomical differences (Fig. 2 and 3). These eight locations were indented by transverse lines of 10 indentations, with a step size of 50 µm (Fig. 2). The indentation lines (500 µm length) were centered to the embryo midline so that on every rostrocaudal position, five regions of interest were measured: lateral mesoderm (regions 1 and 5), paraxial mesoderm (regions 2 and 4) and the midline (region 3) (Fig. 3). A total of 20 embryos were explanted and examined, of which 12 embryos were used for the experiments. Other embryos were either damaged or detached during the measurements. All stiffness values are reported as mean ± SEM.

## Data availability

All raw and processed data of this study are available from the corresponding author on reasonable request.

## Supplementary figures

**Figure S1.**
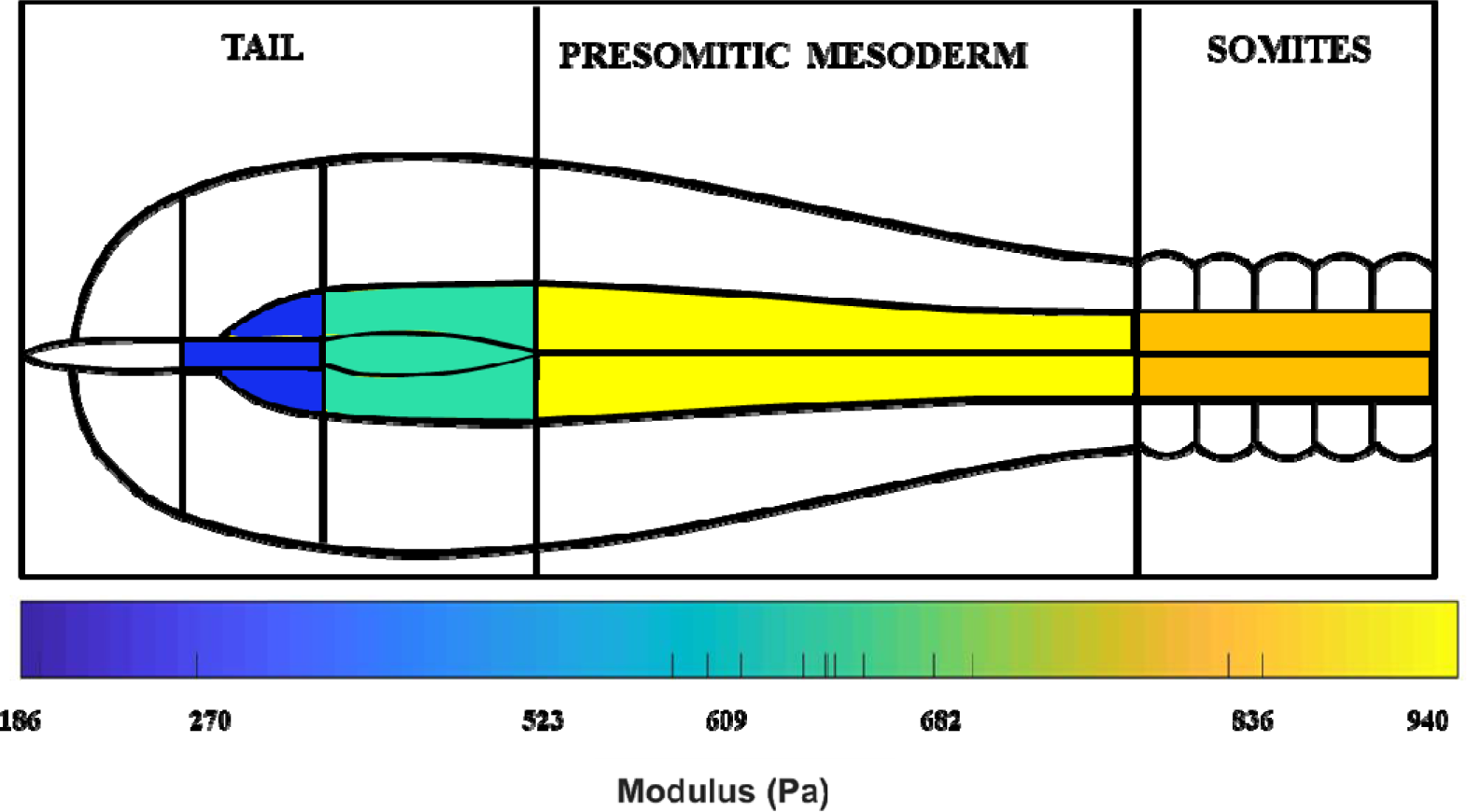
Averaged stiffness () gradient from tail to somitic region.

**Figure S2.**
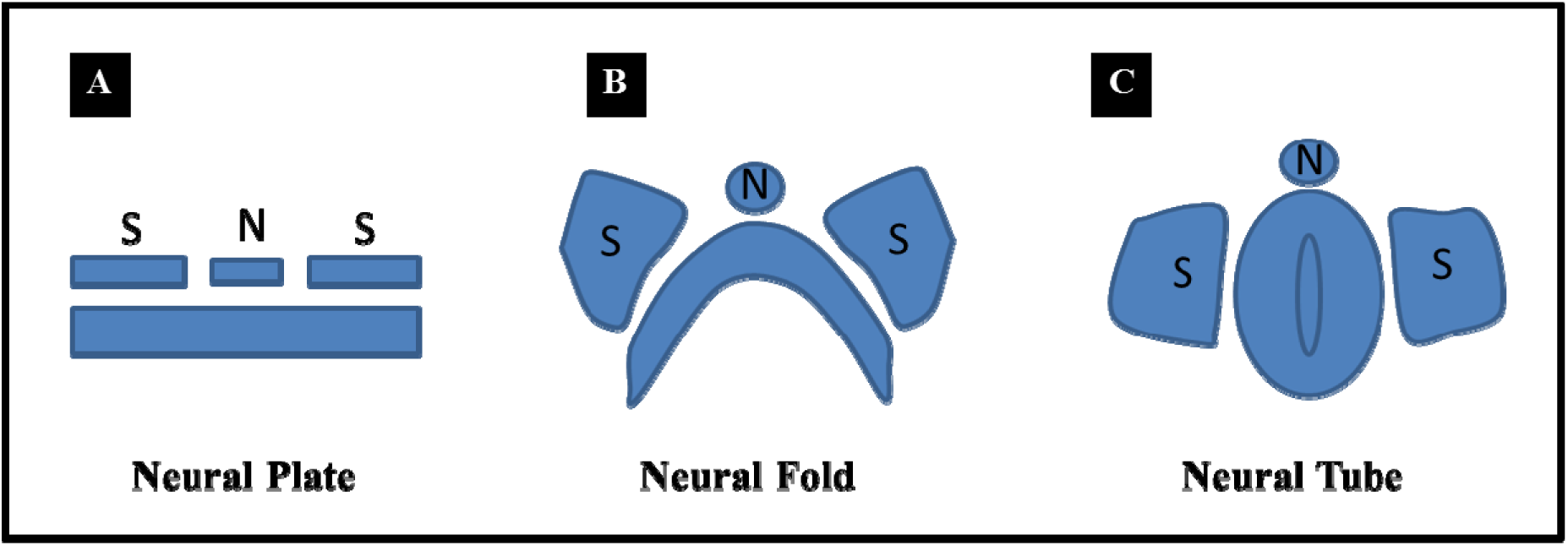
Development of the embryonic spinal cord. (A)The neural plate is generated as a columnar epithelium, then it starts to fold into a neural fold (B), and then the fusion of the neural folds forms the neural tube (C). Abbreviations: (S) somite, (N) notochord. Drawing not to scale.

**Figure S3.**
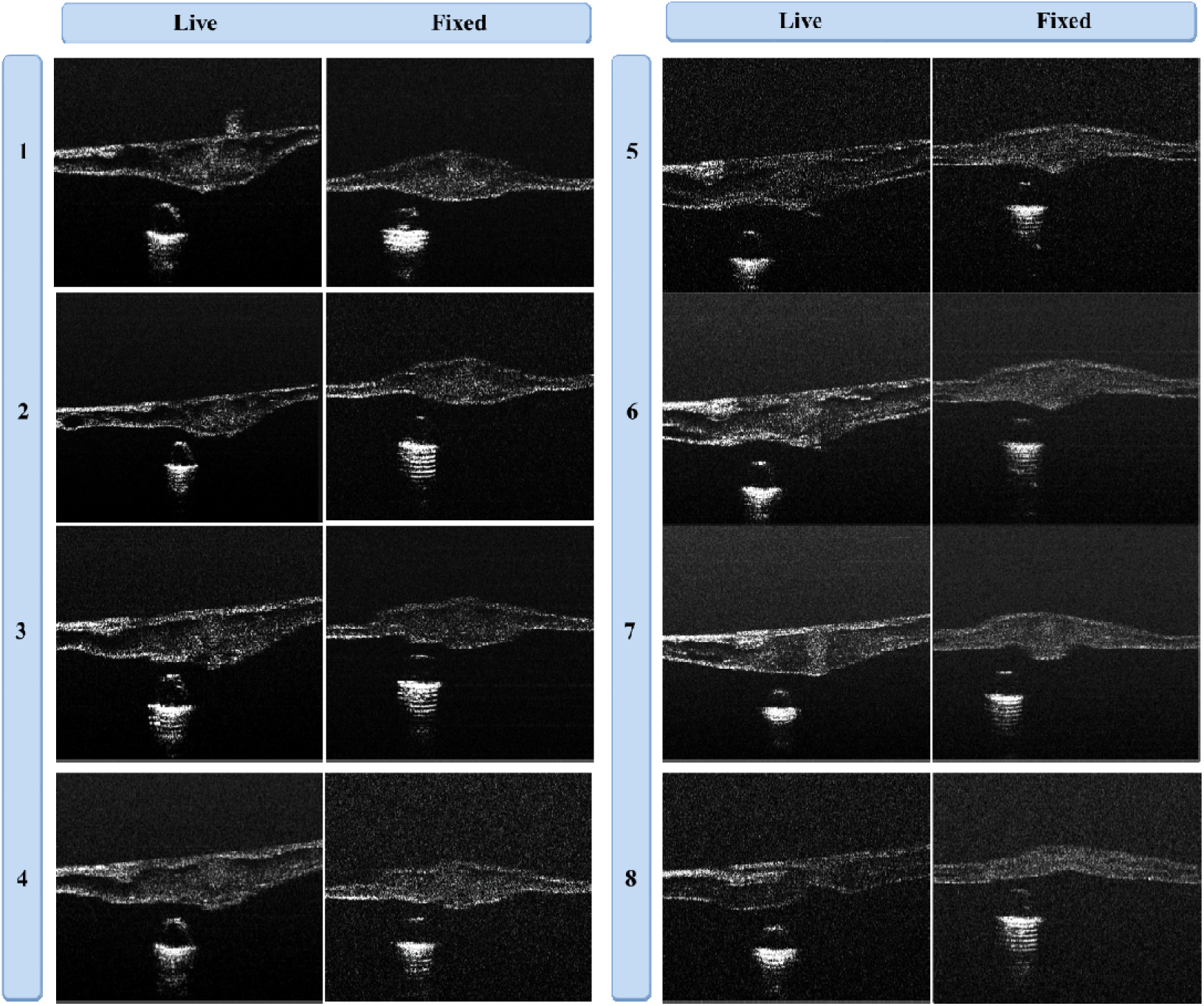
Transverse OCT images of fix vs. live embryo. Images of eight indentation lines on the same live and fixed embryo (the embryo has been imaged live, fixed for 2 hours and then imaged again at approximately the same locations).

